# GAPiM: Discovering Genetic Variations on a Real Processing-in-Memory System

**DOI:** 10.1101/2023.07.26.550623

**Authors:** Naomie Abecassis, Juan Gómez-Luna, Onur Mutlu, Ran Ginosar, Aphélie Moisson-Franckhauser, Leonid Yavits

## Abstract

**Variant calling is a fundamental stage in genome analysis that identifies mutations (variations) in a sequenced genome relative to a known reference genome. Pair-HMM is a key part of the variant calling algorithm and its most compute-intensive part. In recent years, Processing-in-Memory (PiM) solutions, which consist of placing compute capabilities near/inside memory, have been proposed to speed up the genome analysis pipeline. We implement the Pair-HMM algorithm on a commercial PiM platform developed by UPMEM. We modify the Pair-HMM algorithm to make it more suitable for PiM execution with acceptable loss of accuracy. We evaluate our implementation on single chromosomes and whole genome sequencing datasets, demonstrating up to 2x speedup compared to existing CPU accelerations and up to 3x speedup compared to FPGA accelerations.**

## I. Introduction

The completion of the Human Genome Project in 2003 and the advent of next-generation sequencing (NGS) technology in the early 2000s marked the beginning of a new era in DNA sequencing [1,2], enabling faster and more efficient sequencing of large amounts of DNA, and driving the growth in the field of genomics. Over the years, the progress made in the field has led to a rapid increase in the amount of available genomic data and a decrease in the cost of sequencing.

Genomic data is analyzed to understand the genetic information carried by a DNA molecule and to identify DNA alterations that can cause diseases, such as genetic disease and cancer. The process of identifying variations in the sequenced data compared to a reference genome is known as variant calling and can take up to 40% of the total time spent in the genome sequencing pipeline [4]. Variant calling is an essential step in DNA sequencing pipelines because it enables the identification of differences, or variations, between the DNA sequence being analyzed and a reference genome or other sequences. These variations can include single nucleotide polymorphisms (SNPs, i.e., a DNA sequence variation that occurs when a single nucleotide (adenine, thymine, cytosine, or guanine) in the genome sequence is altered), small insertions or deletions (indels^2^), or larger structural variations such as copy number variations (CNVs, i.e., refer to the genetic trait involving the number of copies of a particular gene present in the genome of an individual) or genomic rearrangements.

Variant calling is important for a variety of applications, including disease diagnosis [18,20], personalized medicine, and understanding evolutionary relationships between organisms [19]. By identifying genetic variations, variant calling can help researchers and clinicians identify disease-causing mutations, develop targeted therapies, and better understand the genetic basis of disease. In addition, variant calling can be used to study the genetic diversity of populations or track the spread of infectious diseases.

With the increased need for genomic data and continuous advancements in technology, it is predicted that the amount of genomic data generated will continue to rise [5]. Most of the computational algorithms developed for genome analysis are designed for use in a CPU system. The analysis of the sequenced data requires intensive data movement between DRAM and processing elements which are expensive, both in terms of time and energy [6,7,8,9]. In addition, the variant calling task is a highly parallelizable task as different regions in the genome can be analyzed independently of the others. While CPUs, FPGAs and GPUs are commonly used in variant calling to take advantage of parallel processing [30,31,32,33,35,36], the speedup achieved by these hardware architectures is limited as explained in the evaluation section, emphasizing the need for a new architecture paradigm.

In this paper we focus on Processing-in-Memory computer architecture paradigm whose main purpose is alleviating the *memory wall*, which limits the performance and energy inefficiency of a conventional von Neumann architecture for data intensive applications such as genome analysis. Processing in Memory (PiM) architecture is a promising solution for genomic data analysis considering the expansion of the available data [10,11,12,13,14,15]. PiM architectures can solve data access bottleneck and lower the energy consumption.

Pair-HMM is the most time-consuming part of the variant caller in the GATK pipeline. Thus, we investigate the PiM architecture as a potential solution of acceleration. PiM architecture provides high bandwidth and low latency memory access. Additionally, PiM provides a large amount of compute parallelism and enables a throughput that scales with memory size.

In this work, we show that a PiM architecture can efficiently implement an end-to-end solution for genomic analysis, even when certain components of the genome analysis pipeline are not highly data intensive.

The main contributions of this work are as follows:

1. We rethink and adapt the Pair-HMM algorithm to make it suitable for a real Processing-in-Memory architecture.
2. We implement the Pair-HMM algorithm on a commercial PiM system with more than 2500 PiM cores (DPUs).
3. We comparatively evaluate the PiM Pair-HMM implementation and demonstrate up to 2x time performance improvement compared to high-performance CPU and up to 3x performance improvement compared to SOTA FPGA solutions.

## II. Background

### A. The Genome Analysis ToolKit (GATK)

GATK (Genome Analysis Toolkit) is a collection of tools developed by the Broad Institute to identify SNPs and indels in genomic data [16]. SNPs are the most common sort of genetic variation among human beings [17]. SNPs and indels play an important role in the study of human cancer genes [18,19,20]. The GATK best practices pipeline is one of the most widely used for variant calling, due to its high accuracy [21]. This pipeline is composed of three essential parts. The first one is data pre-processing, the second is variant discovery, and the third is variant evaluation. GATK is an open-source software available on GitHub [22].

### B. HaplotypeCaller algorithm

The second part of the GATK pipeline, variant discovery, is implemented by the HaplotypeCaller algorithm. This algorithm gets as inputs a processed BAM file (a set of aligned DNA reads) and a reference sequence file, and outputs a VCF file (variant calling file). The HaplotypeCaller algorithm can take up to 35% of the complete genome analysis pipeline time as reported in [23]. The HaplotypeCaller algorithm has four main steps:

- *Active region determination* based on the presence of a sufficient number of variations that exceed a predetermined threshold [16]. Active regions, i.e., with sufficient variations, are the target of the next three steps.
- *De-Bruijn like graph assembly* is used to assemble the reads of the region and determine a list of haplotypes [24]. A haplotype is a sequence that represents a certain combination of alleles (alternative gene forms) present in a region. Then the Smith-Waterman algorithm is used to align each haplotype to the reference haplotype to identify potential genetic variations.
- *Pair-HMM forward algorithm* is used to perform a pairwise alignment of each read against each haplotype [25]. The algorithm calculates the likelihood of each haplotype given each read by summing the likelihoods of all possible alignments of the read to the haplotype. This likelihood reflects the probability that the read and the haplotype are related, based on their sequence similarity.
- *Genotype sampling* estimates the genotypes (combination of alleles) at each variant site in the region using Bayes’ theorem in combination with the previously obtained likelihoods.

### C. Algorithm analysis

The GATK HaplotypeCaller software includes a feature that tracks the time required to complete the Smith-Waterman and the Pair-HMM steps of the workflow. Additionally, we used profiling tools [55,56] and present the time repartition of the program in *Figure 1* (running on an Intel Core I7-5820K x86_64, 6 cores (12 threads), 3.30GHz, 64GB). The figure represents the average relative execution time for chromosomes 1-3 of sample NA12878 (commonly used sample from a female individual of European descent). As observed, the Pair-HMM component of the algorithm accounts for a significant portion of 49% of the total time, prompting our decision to accelerate this section.

**Figure 1.**
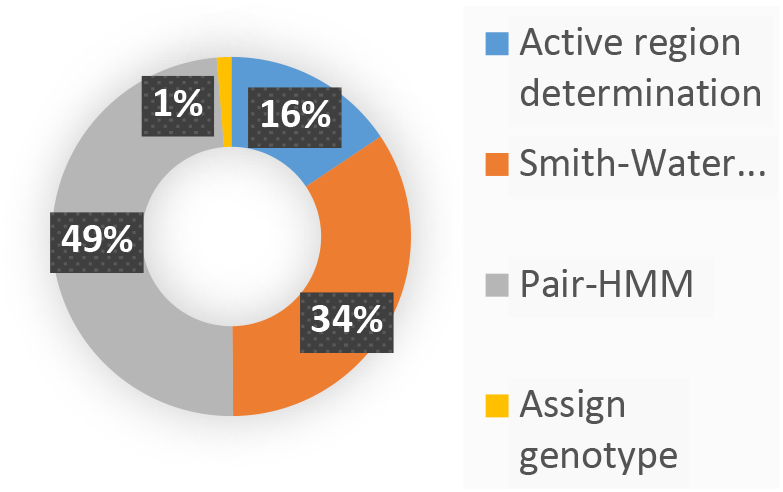
Execution time breakdown of GATK HaplotypeCaller.

Moreover, we check the memory boundedness of the HaplotypeCaller algorithm by using hardware performance counters [52]. As reported in [52], the memory boundedness of the program, which is a component of the non-execution of the CPU (*cycle_activity.cycles_no_execute*), can be computed by the following formula:

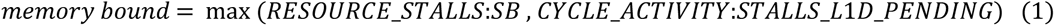

We present in the following table the results of this analysis for a single chromosome (chromosome 1 randomly chosen) and for a WGS running. As can be observed, the memory boundedness of the HaplotypeCaller algorithm is low, which means that memory access is unlikely to become a bottleneck of a HaplotypeCaller. The intuitive conclusion is that PiM would not be the most efficient solution for HaplotypeCaller. Nevertheless, there are three main advantages to PiM architecture even this non-memory-bound program can benefit from:

1. PiM architectures have a large number of execution units that provide large amounts of parallelism.
2. PiM compute throughput scales with memory capacity.
3. PiM systems are not limited in terms of memory capacity (or not as much) as other accelerators (e.g., GPUs or FPGAs with HBM memory).

**Table 1.**
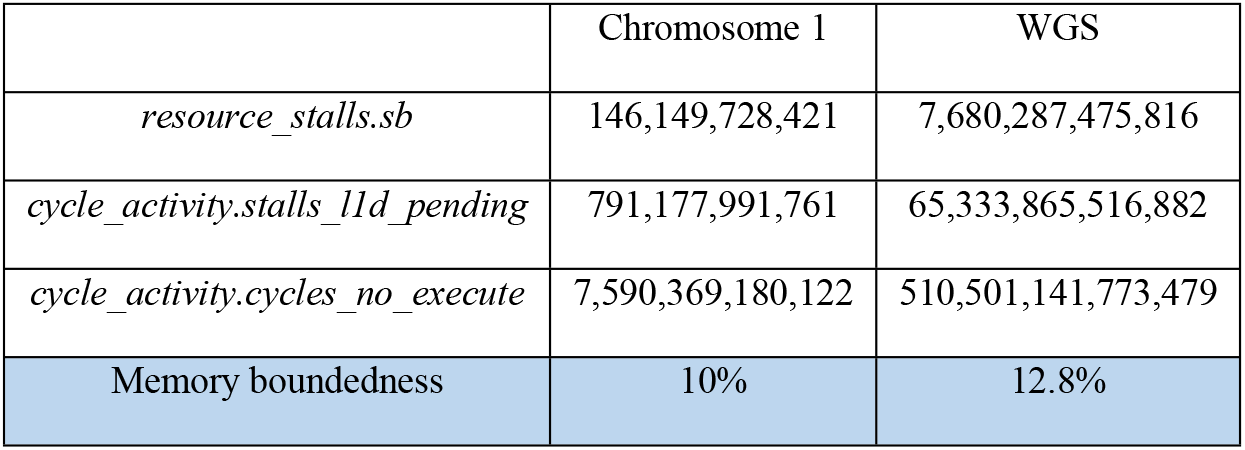
Memory boundedness of HaplotypeCaller algorithm.

### D. Pair-HMM forward algorithm

In this work we focus on the Pair-HMM step which is the most resource-intensive part of the HaplotypeCaller algorithm. In this part we compute the likelihood of each read against each haplotype.

The likelihood calculation is based on a pairwise alignment using a Hidden Markov Model (represented by a state machine) and the qualities of the read DNA bases (qualities produced by a sequencer). Given a DNA read R of length *n* and a haplotype H of length *m* we define the transitions of the state machine as the probabilities of the possible changes in state that occur during the process of aligning the read to the haplotype. In other words, the state machine represents the different states that the alignment can go through, such as a match (state M), an insertion (state I) or a deletion (state D) in the read. The transitions between the states are defined by the following probabilities:

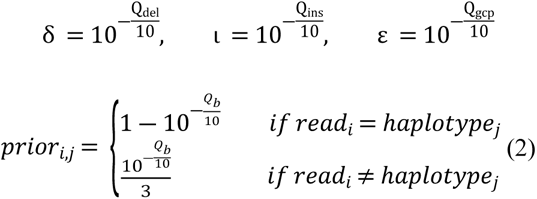

Currently, Q_del_, Q_ins_ and Q_gcp_ (respectively the deletion, insertion, and gap continuation qualities) are included in the PHMM model but are not produced by sequencers and thus default values determined by GATK [22] are used for each base. However, Q_b_ is referred to as the quality of each base in the read and is provided by the sequencer machine.

The finite state machine of *Figure 2* models the transitions between the three different states (M, the match state, I, the insertion state and D, the deletion state).

**Figure 2.**
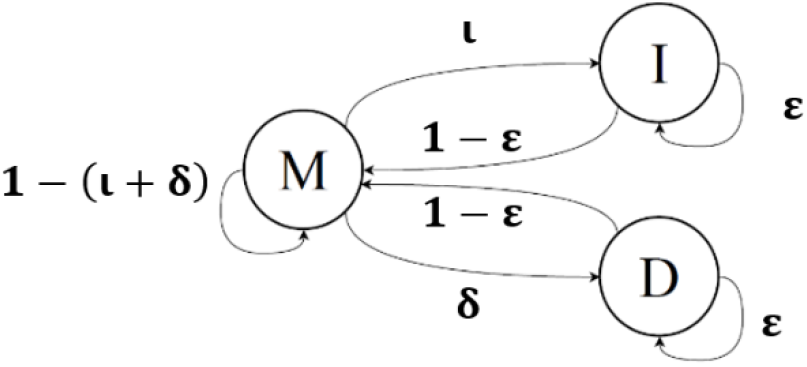
Finite state machine model for HaplotypeCaller.

Using this state machine with dynamic programming, the algorithm fills up three matrices, M, I and D as shown in Equations *(3;1)* and *(3;2)*. Each element *i*,*j* of the matrix represents the total likelihood of all paths from the beginning of the read to position *i* such that the path ends with the corresponding state (match state for matrix M, deletion state for matrix D, and insertion state for matrix I).

*Initialization:* ∀ 0 ≤ *i* < *n*, 0 ≤ *j* < *m*,

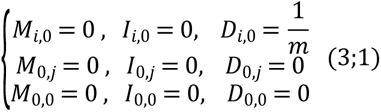

*Recursion step:* ∀ 1 ≤ *i* ≤ *n*, 1 ≤ *j* ≤ *m*,

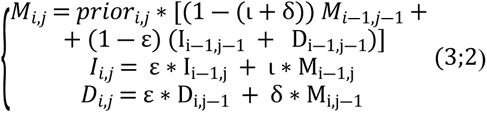

*Termination step:*

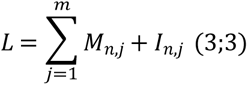

The terminations step computes the final likelihood for the current read-haplotype.

## III. Related work

Genome analysis requires high-performance computing to process and store large and continuously growing genomic datasets. Several processing in memory (PiM) solutions have been proposed to accelerate various parts of genome analysis pipeline. For example, the development of UpVC [13] on the UPMEM architecture has sped up the read alignment and variant calling process and yielded a time and energy improvement compared to GPU or FPGA implementations, using a simpler and less accurate algorithm than PHMM. Other works have focused on accelerating the read alignment part of the pipeline by using in-memory databases [26,27], or by using resistive content addressable memory (BioSEAL [11]). RASSA [28] accelerates read pre-alignment using approximate search-capable resistive content addressable memory. GRIM filter [29] integrates computation within a logic layer optimized to exploit 3D stacked memory layers to filter seed location. More recently GenStore [51] has implemented filters for in-memory reads for genome analysis tasks, and research conducted by [15] demonstrates that the wavefront algorithm achieves greater read alignment throughput on PiM architecture. In contrast to the majority of prior research, it is crucial to emphasize that our work is conducted on a real Processing-in-Memory architecture.

GAPiM is the first GATK pipeline implementation on a commercial PiM platform. To our knowledge, GATK has never been implemented on an actual PIM platform before. Typical GATK accelerators are based on CPU, FPGA, and GPU. Most of these accelerations implement the Pair-HMM step. CPU acceleration solutions use a combination of SIMD extensions such as AVX and AVX-512 and multithreading [30]. FPGA-based solutions mainly focus on accelerating the floating-point calculations of Pair-HMM. Most FPGA solutions use systolic arrays [31,32,33]. Another solution for FPGA is to create a specialized unit to execute the Pair-HMM command and replicate it to provide efficient parallelization [34]. But these FPGA solutions suffer from limited on-chip memory. Our solution, GAPiM, alleviates the memory wall which may limit the CPU and FPGA implementations and hence enables better performance and scalability. We believe PiM approaches are also beneficial to investigate for emerging deep learning based variant calling tools [58]. We leave the investigation of the evaluation, design, and analysis of such tools to future work.

GPU accelerations are also used since the Pair-HMM algorithm is easily parallelizable, focusing on intra-task and inter-task parallelism [35] and allowing to retrieve the optimal alignment [36]. Recently, an ASIC that replaces floating point multiplication by 20-bit addition in log domain employs bound checks to maintain correctness has been proposed in [37]. GAPiM provides a better scalability and flexibility when compared with GPU or ASIC implementations.

## IV. UPMEM PiM architecture

### A. UPMEM PiM chip

In this work we utilize the PiM platform developed by UPMEM [39]. It is the first commercially available PiM solutions. It has been shown to be effective in speeding up memory-intensive workloads [13,40,41]. UPMEM platform is a standard DDR4-2400 DIMM with several PiM chips, each one composed of 8 parallel processors, called DPUs (DRAM Processing Units). Although this paper we refer to DPUs as PiM cores. The PiM cores are operated by a host CPU that sends the program to be executed and collects the processing results when PiM cores complete their processing tasks [39,42,43].

### B. Chip level organization

Within each memory chip, there are 8 asynchronous PiM cores that operate independently of one another. Each PiM core is associated with a 64MB DRAM bank called MRAM (Main RAM) that can also be accessed by the host CPU. Different PiM cores cannot communicate with each other directly. The host CPU can transfer data to/from MRAM banks of individual PiM cores. This design allows each computing unit to efficiently process its own fragment of the dataset. Calculations are performed in-situ, within each unit at a memory bandwidth of almost 1GB/s.

**Figure 3.**
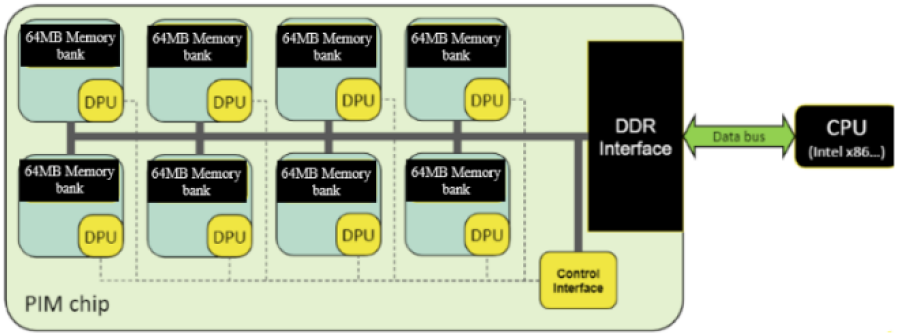
UPMEM PiM architecture [39].

### C. The PiM core

A PiM core is a 32-bit RISC-based processor (with 64-bit capabilities) running at 400MHz. It is a multithreaded processor with up to 24 hardware threads, called PiM threads. The PiM core has a pipeline depth of 14 stages, however, only the last three stages of the pipeline can execute in parallel with stages of the next instruction in the same thread. Therefore, instructions from the same thread must be dispatched 11 cycles apart, requiring at least 11 threads to fully utilize the pipeline [39,42,43]. Each PiM core has an exclusive access to a 24KB IRAM (Instruction RAM), and to a 64KB WRAM (Working RAM) shared by all the threads, as well as the 64MB MRAM as illustrated in *Figure 4*. The WRAM has a lower access latency than MRAM and can be accessible in one cycle. To allow several threads to access a same resource we use a mutual exclusion (mutex) mechanism, i.e., a synchronization object is used to control access to a shared resource and ensure that only one PiM thread can access that resource at a time, avoiding conflicts and data inconsistencies.

**Figure 4.**
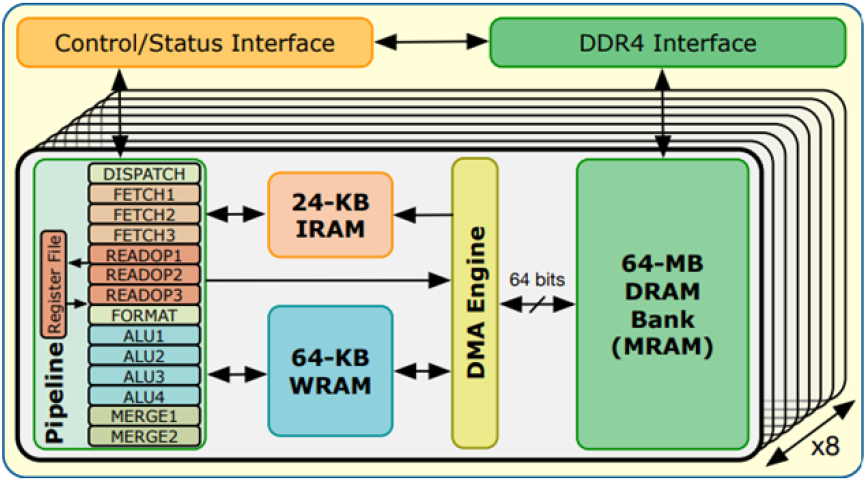
UPMEM chip diagram (reproduced from [38])

In this setup, programmers explicitly transfer data between MRAM and WRAM using DMA calls which must be 8-byte aligned and execute load and store operations on WRAM.

## V. Adapting HaplotypeCaller algo-rithm to UPMEM PiM architecture

We modified the Pair-HMM algorithm to enable its efficient implementation on UPMEM PiM architecture while optimizing the hardware utilization.

### A. Proposed log domain fixed point model

The need to redesign the Pair-HMM for PiM core stems from two observations, as follows: first, multiplications in PiM core are much slower than additions (approximately 10 MOPS for multiplication vs. 60 MOPS for 32-bit integer addition) [40]. Second, floating point operations are significantly less efficient on PiM cores compared to integer operations (approximately 5 MFLOPS for additions) [40]. This is because the PiM core does not implement floating point ALUs and these operations are emulated by the UPMEM runtime library in software [44, 45].

The Pair-HMM algorithm requires a high number of multiplications (Equations *(3;1)* and *(3;2)*) and uses floating point calculations (see priors and transition probabilities). These calculations are significantly slower than integer addition/subtraction on the wimpy PiM cores: multiplication takes six times longer and floating-point addition takes twelve times longer than integer addition. To overcome this limitation, we first convert all equations to log domain, replacing multiplications with additions. For example, the first equation of *Equation (3;2)* is transformed as follows, computing the value of log (*M*_*i*,*j*_) rather than *M*_*i*,*j*_:

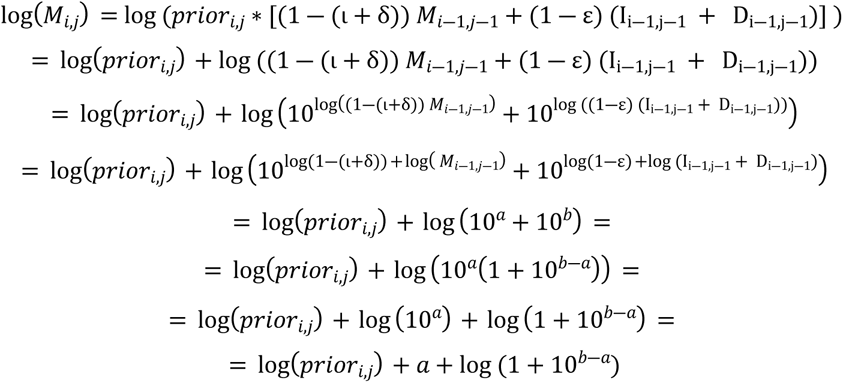

Where *a* = log(1 ― (ι + δ)) + log(*M*_*i*―1,*j*―1_) and *b* = log(1 ― ε) + log (I_i―1,j―1_ + D_i―1,j―1_).

Several methods for PiM execution of log and other transcendental functions have been proposed **Error! Reference source not found.**. To implement log-domain additions, we use a look-up table (LUT) to quickly calculate the value of log(1+10^X^). This allows us to compute the value of the logarithmic *sum* (*log*(*x* + *y*)) according to the above equations. Such LUTs are implemented in PiM core memory prior to the operation. The log values of constants and known-in-advance initial values are precalculated by the host CPU and stored in the PiM core memory prior to the operation.

Our second contribution is replacing the floating-point calculations by fixed point ones. This reduces the number of slow multiplications and eliminates the need for emulated floating-point operations, considerably improving the performance and efficiency of the algorithm implementation on PiM cores. We modify the Pair-HMM recursion step as shown in *Equation (4)* below, where + _*F*_ represents a fixed-point regular addition and + _*L*_ represents a logarithmic addition (i.e., log(a+b)) performed using the LUT approach.

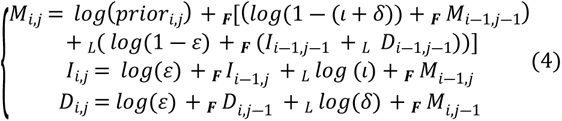

### B. Accuracy analysis

Replacing the floating-point operations by fixed-point ones affects the computation precision and therefore, impacts the accuracy of the algorithm (i.e., the ability of the algorithm to correctly identify all the variants). We investigate this accuracy impact by comparing our modified GATK implementation with the original GATK HaplotypeCaller algorithm.

We use two samples from the International Genome Sample Resource (IGSR) [46], NA19685 and NA12878, for each one WES (Whole Exome Sequencing, i.e., protein-coding regions of genes in the genome [47]) with 30x coverage) and WGS (Whole Genome Sequencing) with low coverage. These genomes are aligned to the Hg38 reference and the GATK best practices pipeline is applied on each one.

We optimize the fixed-point format of our calculations using the Matlab Fixed-Point Designer tool [48]. This tool suggests the optimal fixed-point configuration for the variables involved, ensuring accurate and reliable results. It enables us to select the appropriate wordlength for each variable, minimizing the accuracy loss in the Pair-HMM execution.

To compare the performances of the variant callers we first define the following:

- True positive (TP): variants found by both the original GATK and by fixed-point precision GATK.
- False Negative (FN): variants found by original GATK but not found by fixed-point precision GATK.
- False Positive (FP): non-existing variants found by fixed-point precision GATK.

The metrics of precision (P) and sensitivity (S) are defined as follow, the reference being the original GATK program:

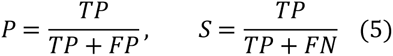

We use the *vcfeval* tool [50] to compute the different metrics testing four samples (Whole Exome Sequencing (WES) and Whole Genome Sequencing (WGS)) of two different subjects (NA19685 and NA12878). Our analysis shows that fixed-point utilization affects the VCF output, as can be seen in *Table 2* below. However, the extent to which this impact is acceptable or not may depend on application requirements and should be evaluated accordingly.

**Table 2.** Fixed-point impact on sensitivity and precision.

| Sample | NA19685<br>WES | NA19685<br>WGS | NA19685<br>CHR18 | NA12878<br>WES |
| --- | --- | --- | --- | --- |
| Coverage | High | Low | High | High |
| Bits | 32 | 32 | 12 | 12 |
| Decimal bits | 25 | 25 | 7 | 8 |
| P | 0.9998 | 0.9996 | 0.9717 | 0.9993 |
| S | 0.9998 | 0.9996 | 0.9996 | 0.9996 |

### C. PiM core initialization

We pre-calculate the log of the initial values and constants in the host CPU since the log function does not exist in the PiM core runtime library. These log values are then transferred in a fixed-point format directly to the PiM core. For instance, the initialization value for *D*_*i*,0_ is sent to the PiM core as 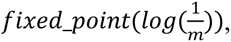 and the transition values are sent as the fixed point of their log values.

## VI. Implementation on PiM cores

### A. CPU-PiM data transfers

The data of all the regions assigned to a particular PiM core is transferred to the PiM core’s MRAM. It includes the reads of the regions and their auxiliary information (read bases, length, and log-domain transitions probabilities), as well as the haplotypes of the regions and their information (read sequence, length and log-domain initial value (*fixed_point(log(1/m))* as detailed in Section V.C.)).

### B. PiM kernel

Each PiM thread is assigned a read; then each PiM thread calculates the likelihood of such read against each haplotype in the region. Whenever a read from a new region is assigned to the PiM core, the haplotypes for that region are transferred to the WRAM and remain until the processing for the region is completed. This approach allows us to effectively exploit data reuse, as the haplotypes can be accessed multiple times during the region processing.

Each running PiM thread fetches (function *reserve_read* in *Algorithm 1*) a read from a pool of reads (either at the start of the program or when it completes handling its current read) and then proceeds to transfer the relevant data associated with that read to the WRAM to compute the likelihood result of the associated read against each haplotype of the region. To manage the read allocation to PiM threads, we maintain a global counter that is shared among all PiM threads, and use a mutex to prevent simultaneous allocation of reads to multiple PiM threads, as explained above (part IV.C).

A simplified pseudo-code run by each PiM thread is presented in Algorithm 1.

**Algorithm 1.**
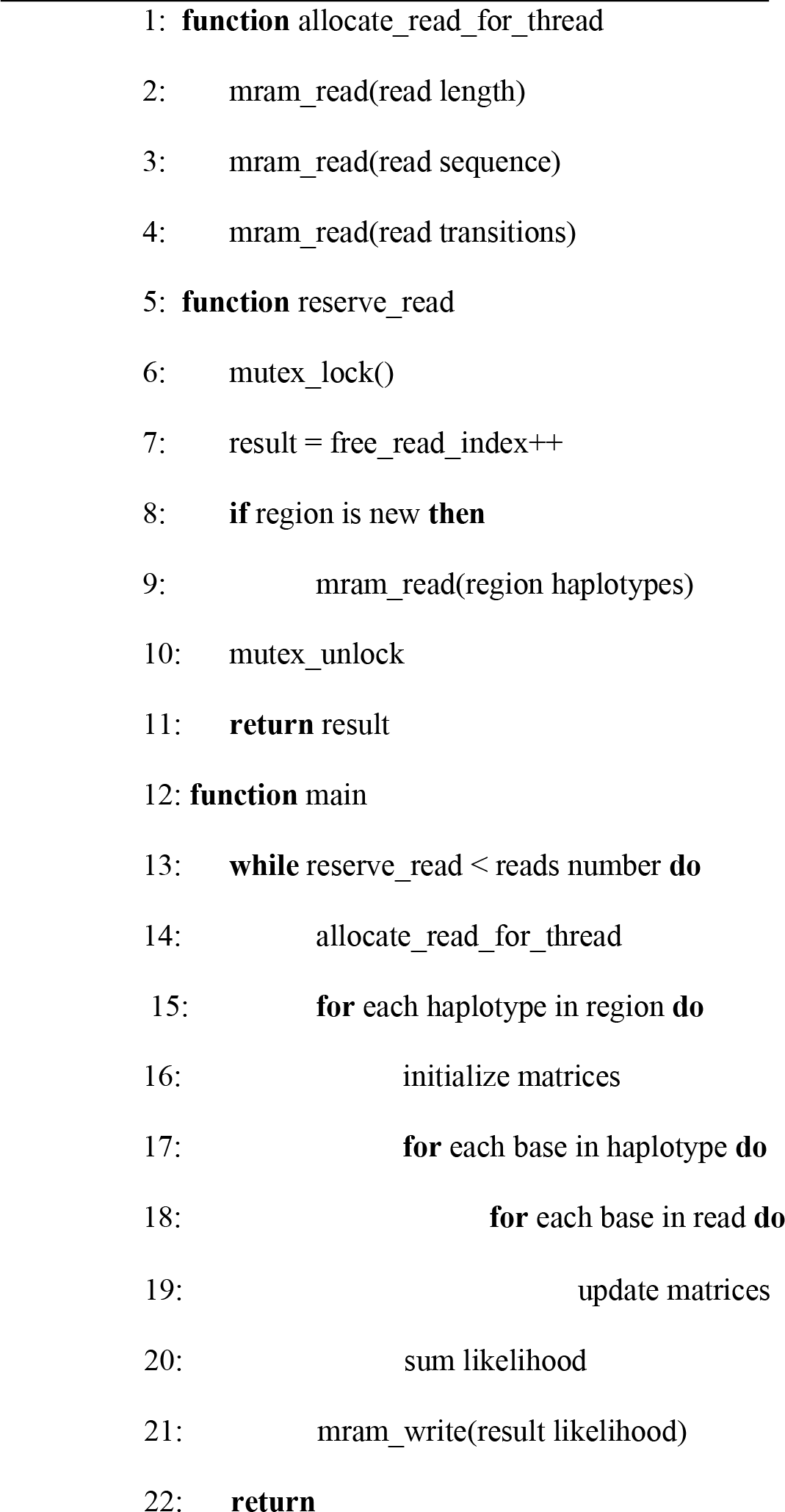
Pseudo code for PiM core.

### C. Efficient handling of on-chip memory

To optimize the memory usage, the haplotypes present in the WRAM (that have been transferred from the MRAM) are stored in a circular buffer. The utilization of the circular buffer allows for a continuous reuse of memory by replacing the oldest data with new data, preventing wastage, and making efficient use of the available memory. Next, we transfer the read information from the MRAM to the WRAM, including the length, sequence, and transitions probabilities (respectively lines 2,3,4 in *Algorithm 1*).

In order to fill the matrices efficiently and optimize memory usage, we only keep two lines of each matrix (for each PiM thread). This is because, as presented in Equation (2), the value in each matrix cell depends only on its neighbors as illustrated in *Figure 5*. The intermediate likelihood result for each read-haplotype pair is stored in the WRAM. After completing the pairwise alignment of the read and the haplotype, the resulting likelihood is written to the MRAM memory in a result matrix (line 21 in *Algorithm 1*). In order to maximize the bandwidth of the WRAM-MRAM transfer, we coalesce writes from several PiM threads.

**Figure 5.**
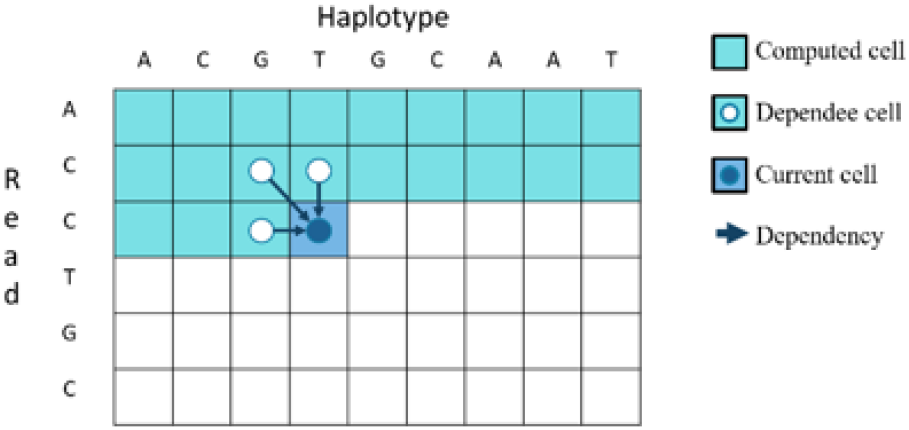
Cell computation dependencies in the matrix.

### D. Load balancing over PiM cores

To optimize the performance of the parallel PiM cores, it is important to ensure that the workload is evenly distributed among them. One alternative approach to the method we employ in this work involves dividing genome regions using a SNP database, as reported in [38], assuming that the number of variants in the sample is comparable to the number of variants in the database.

Our approach is to allocate several regions to each PiM core based on their complexity, which is determined by the total length of reads and haplotypes in the region. The load balancing is done by the host CPU that evaluates the complexity of each region and potentially divides regions into smaller fragments to meet a specified complexity target.

### E. Asynchronous implementation

As presented in Section V, the pair-HMM algorithm requires redesign to run efficiently on PiM cores, including pre-processing and post processing steps running on the host CPU, such as splitting the regions. PiM cores can be launched asynchronously, i.e., we can launch them by rank of 64 PiM cores and each rank can be launched independantly of the others. We leverage this to minimize the load imbalance among PiM cores ranks. To efficiently utilize available ressources and optimize PiM cores work we construct a pipeline as illustrated in *Figure 6* below. To link between the different stages of this pipeline we use buffer queues (one input queue and one output queue). This pipeline is executed on the host CPU and contains three stages:

- *Pre-processing stage* handled by one single thread that writes to the input queue.
- *PiM cores populating and launching stage*. In this stage, a thread is created for each PiM core rank and reads from the input queue to transfer data to PiM cores and writes to the output queue the final result produced by the PiM cores.
- *Post processing stage* handled by a single thread that reads from the output queue and writes the result to an output file.

**Figure 6.**
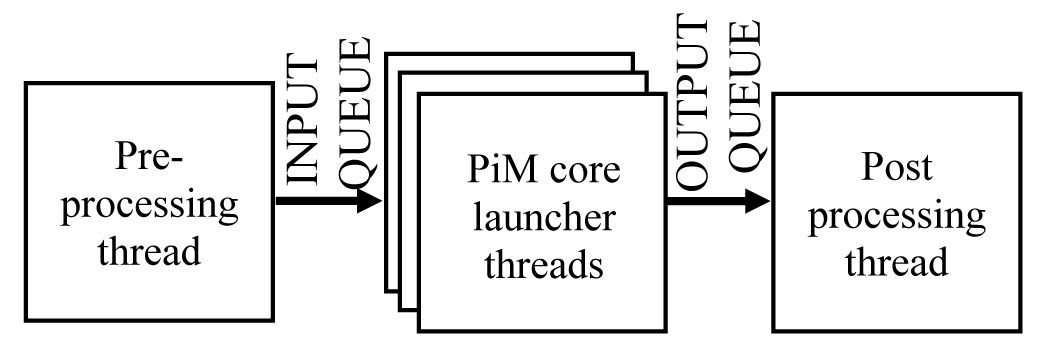
Host CPU pipeline stages.

By constructing the pipeline in this manner, we are able to achieve latency hiding during the pre-processing, data transfer, and post-processing steps. This is because the time spent on PiM core computation is much greater than the time spent on these tasks. Thus, when computing the running time on future architectures (as discussed in VII) we disregard the impact of non PiM core time, as explained in [53].

The profiler tool provided by UPMEM [59] illustrates the pipeline parallelization as shown in *Figure 7,* with two levels of parallelism: the first is inter-rank parallelism, meaning that several ranks run in parallel (green segments represents PiM core time), and the second is thread parallelism on the host side, meaning that pre-processing task (blue segments), data transfer task (red segments), and post-processing tasks (orange segments) are performed concurrently.

**Figure 7.**
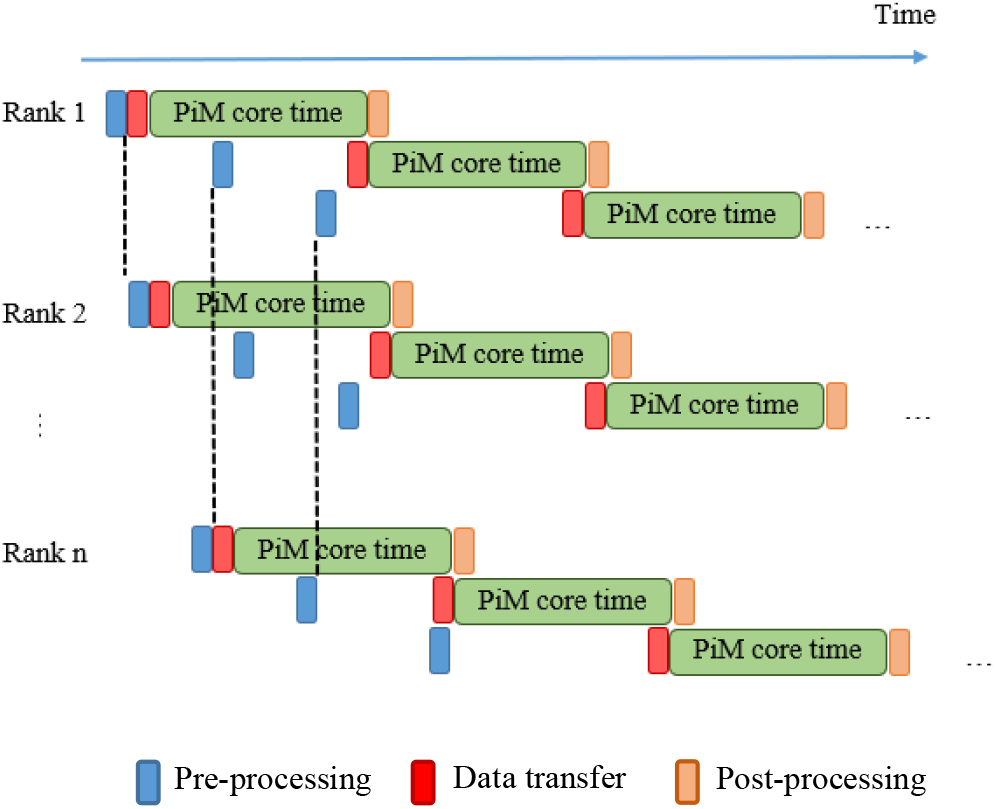
Time diagram of the system pipeline.

## VII. Evaluation

### A. Evaluation Methodology

The algorithm is implemented on servers provided by UPMEM. The architecture is as follows:

- Host: Intel Xeon Silver 4215 CPU, 2.5GHz
- 2560 PiM cores divided into 40 ranks.
- Each PiM core operates at a frequency of 400MHz.

The benchmarks used in this paper are obtained using a publicly available Whole Genome Sequencing dataset of NA12878 from the International Genome Sample Ressource (IGSR) [46]. We use real datasets in order to ensure transparency and accurately reflect the performance of the Pair-HMM algorithm in real-world scenarios. We present the results for chromosomes 1, 2 and 18 as well as for WGS dataset.

We compare the processing time of our solution with different CPU implementations, including the original Java implementation, the AVX-accelerated version, and the OpenMP multithreaded version (with 4 threads and 8 threads). Additionally, we present the time results of the open-source FPGA implementation from [34], which includes two versions, one with 24 “workers” and the second one with 96 “workers”, where a worker is an acceleration unit (respectively represented as 24wk and 96wk). This Pair-HMM implementation is run on a F1 instance of AWS [54].

### B. Results

The GATK execution time figures are presented in Table 2 below along with the speedup figures, using the original Java implementation [22] of the algorithm as the baseline.

The results indicate that the existing UPMEM architecture (2560 PiM cores operating at 400MHz, indicated by bold numbers) can provide speedups similar to that achieved by CPU or FPGA. For example, the UPMEM platform is able to process chromosome 1 in 117 seconds, which is slightly longer than 100 seconds it took the CPU with OpenMP 4 threads, and slightly faster than the 124 seconds needed for the FPGA implementation. The performance achieved by this existing UPMEM architecture represents up to 10.7x speedup compared to the original Java implementation of the algorithm.

UPMEM has laid plans for continued improvement and innovation of its product offerings, and these results serve as a prediction of what can be expected from these future architectures. We present in *Figure 8* below the performance roadmap [60] of the UPMEM architecture as inspired by [49]. Each percent figure shows the relative improvement of DPU’s frequency in the related time period.

**Figure 8.**
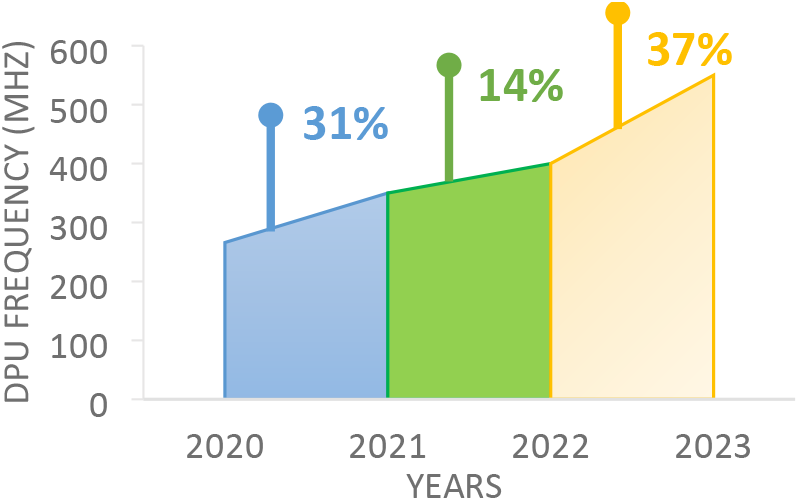
Performance roadmap of UPMEM architecture.

A scaled-up UPMEM platform with 2560 PiM cores running at 550MHz (and above) surpasses both the CPU with OpenMP 4 threads and the FPGA based platforms. This can be seen in the speedup achieved for chromosomes 1 and 2, which are 14.8x and 12.7x respectively compared to 12.5x and 10.6x for CPU (OpenMP 4t) and to 10x and 11.6x for the FPGA speedup respectively. This is a promising result as UPMEM plans on releasing a version of their architecture with PiM cores working at 550MHz in Q2 2023.

**Table 2.** Running time in seconds and acceleration factor of the Pair-HMM algorithm on different architectures.

|  | CHR1 | CHR2 | CHR18 | WGS |
| --- | --- | --- | --- | --- |
| **Java**<br>**(reference)** | 1254.8<br>1x | 638.7<br>1x | 162<br>1x | 15159.6<br>1x |
| **AVX** | 317.1<br>4x | 163<br>3.9x | 40.2<br>4x | 3759<br>4x |
| **OpenMP**<br>**4 threads** | 100.5<br>12.5x | 60<br>10.6x | 15.8<br>10.2x | 1189.4<br>12.7x |
| **OpenMP**<br>**8 threads** | 75.3<br>16.7x | 46.9<br>13.6x | 12.9<br>12.6x | 886.7<br>17x |
| **OpenMP 4t**<br>**FPGA 24wk** | 140.8<br>8.9x | 57.6<br>11x | 12.3<br>13.2x | 1298<br>11.7x |
| **OpenMP 4t**<br>**FPGA 96wk** | 124.4<br>10x | 55.2<br>11.6x | 12.2<br>13.3x | 1200<br>12.6x |
| **UPMEM**<br>**2554 PiM**<br>**cores**<br>**350MHz** | 134<br>9.4x | 79<br>8x | 23<br>7x | 1637<br>9.2x |
| **UPMEM**<br>**2560 PiM**<br>**cores**<br>**400MHz** | **117**<br>**10.7x** | **69**<br>**9.3x** | **20.1**<br>**8x** | **1429**<br>**10.6x** |
| **UPMEM**<br>**2560 PiM**<br>**cores**<br>**550MHz** | 85<br>14.8x | 50.2<br>12.7x | 14.6<br>11.1x | 1039<br>14.6x |
| **UPMEM**<br>**3584 PiM**<br>**cores**<br>**550MHz** | 60.8<br>20.6x | 35.8<br>17.8x | 10.4<br>15.6x | 742.3<br>20.4x |
| **UPMEM**<br>**3584 PiM**<br>**cores**<br>**600MHz** | 55.7<br>22.5x | 32.8<br>19.5x | 9.6<br>16.9x | 680.5<br>22.3x |
| **UPMEM**<br>**5120 PiM**<br>**cores**<br>**600MHz** | 39<br>32.2x | 23<br>27.8x | 6.7<br>24.2x | 476.3<br>31.8x |

*Figure 9* presents (a) the comparison between CPUs execution time (OpenMP four threads and OpenMP eight threads) and PiM cores execution time and (b) the comparison between FPGAs execution time and PiM cores execution time over different architectures for the chromosome 1:

**Figure 9.**
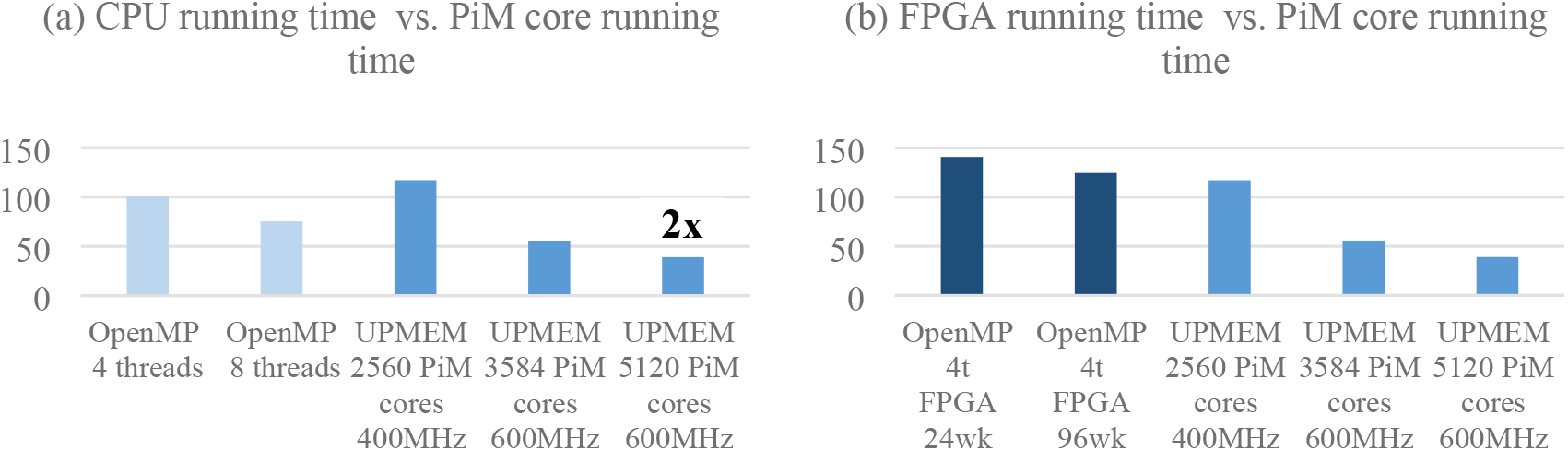
Running time comparison between architectures.

### C. Power consumption and energy

The power consumption of the PiM core depends on its version and clock speed. The actual consumption of a PiM core is not yet available on the current chip version but the PiM core’s TDP (Thermal Design Power, i.e., the measure of the maximum amount of power that the PiM core consumes) is provided by the manufacturer. When running at 350MHz, the TDP of the new version of the PiM core is 112mW, whereas at 400MHz it consumes 220mW [60]. PiM cores that run at 550MHz are designed to consume less energy, and their TDP is expected to be around 200mW.

We compute the overall TDP of the PiM system used for testing, (according to the values provided by the manufacturer [60]) composed of 2560 PiM cores operating at 350MHz. We recall that the UPMEM chip is composed of eight PiM cores and thus this system contains 320 chips.

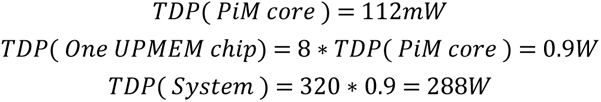

### D. Summary analysis

To summarize this analysis, we compare the speedup achieved by each studied architecture as a function of the number of acceleration units (i.e., the level of parallelism), as presented in *Figure 10* below. The X axis of the graph is the number of acceleration units (as specifically defined in each architecture) and the Y axis of the graph represents the speedup factor for the corresponding accelerator. The PiM UPMEM x-axis represents a projection of performances on future architectures, having as a basis the current existing PiM system (2560 DPUs working at 400MHz).

**Figure 10.**
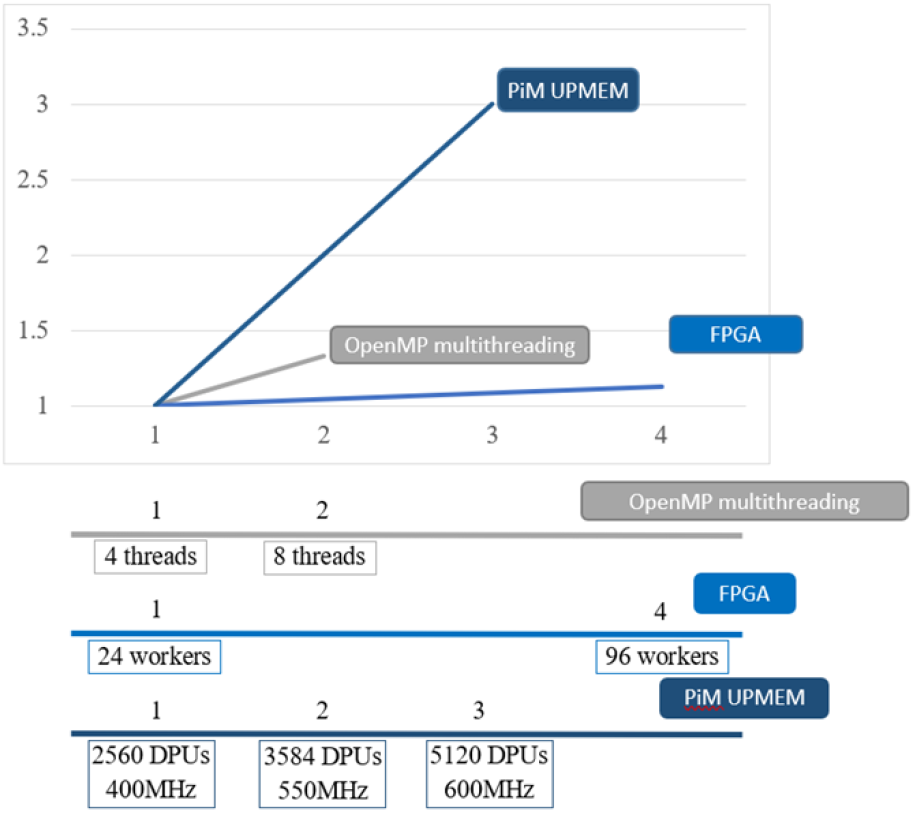
Speedup factor graph for different architectures.

The speedups of OpenMP and FPGA accelerators increase very mildly as the number of acceleration units grows. In contrast, the UPMEM accelerator’s performance scales practically linearly. Indeed, the OpenMP multithreading implementation is limited and suffers from high Cycles Per Instruction (CPI) leading to a low performance increase. Similarly, FPGAs suffer from limited on-chip memory and their high hardware resource consumption makes it impractical to add more workers units.

### E. Discussion

UPMEM PiM platform based Pair-HMM algorithm outperforms state-of-the-art solutions in terms of performance and power consumption. Based on UPMEM performance roadmap, such performance and energy efficiency gaps are expected to grow in the near future. This is because our implementation operates UPMEM DPU at very modest operating frequency which leaves an ample opportunity for further improvements. At the same time, we cannot expect the operating frequency of high-performance CPU and FPGA which are already fully optimized, to improve at similar pace.

Implementing the GATK on UPMEM platforms carries an additional unique advantage. Genome analysis pipeline, which GATK is a major part of, comprises indexing, read mapping, read alignment and in some cases, genome assembly (reference guided or de-novo). Most of these operations are data-intensive and therefore strongly benefit from PiM implementation [10,11,12,13,14,15]. If indexing, read mapping and other data-intensive operations were to be on a PiM platform, while maintaining the conventional CPU-centric (CPU or FPGA based) GATK implementation, it would lead to several significant drawbacks. First, the overall performance would be adversely affected due to the necessity to transfer extremely large amount of sequenced data from PiM platform to a CPU or a FPGA (genomic read file can reach more than 100GB for a single human genome). Second, the total cost of ownership would be significantly higher. Indeed, a heterogeneous solution with two different processing clusters (PiM and FPGA for example) induces higher capital expenditure (CAPEX) due to the cost of the different hardware, as well as higher operational expenditure (OPEX) due to the need for maintenance and increased energy consumption. The ability to implement the entire genome analysis pipeline, from the read mapping to the GATK on the same PiM platform resolves both these inefficiencies. As demonstrated in this paper, to fully realize the benefits of PiM architecture in genome analysis, it is necessary to develop algorithms that are optimized for these architectures, including the parallelization approach, genome region assignment, computing in log domain and so on.

## VIII. Conclusions

This work implements and evaluates Pair-HMM, the main part of the GATK variant calling algorithm and the most computationally intensive part of genome analysis pipeline, on a commercial PiM platform developed by UPMEM. We redesign the original algorithm to optimize it for the UPMEM PiM platform. The suggested modifications include moving the computation to log domain and replacing floating point calculations by fixed point one with very insignificant loss of the variant calling accuracy. To optimize the UPMEM platform utilization and performance, we develop a scheme for optimal allocation of genome fragments to individual PiM cores. This work further demonstrates the potential of PiM architectures as a future solution for end-to-end genome analysis. Despite some parts of the process not being memory-bound, the results show that even complex algorithms such as Pair-HMM can be effectively implemented on PiM architecture and provide speedup over state-of-the-art FPGA or CPU solution.

